## Supplementary figures and images for "Morphological and molecular evidence for range extension and first occurrence of the Japanese seahorse, *Hippocampus mohnikei* (Bleeker 1853) in a bay-estuarine system of Goa, central west coast of India"

### Supplemenatry figure

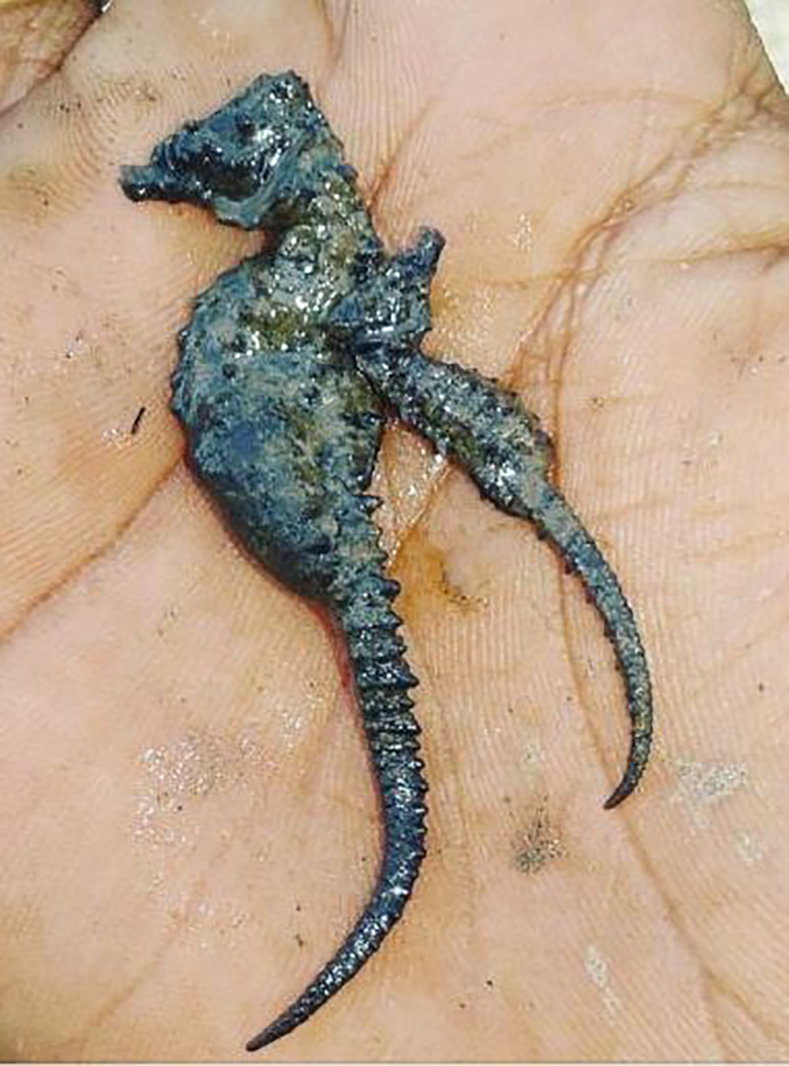
