## Supplementary table 1 for "Morphological and molecular evidence for range extension and first occurrence of the Japanese seahorse, *Hippocampus mohnikei* (Bleeker 1853) in a bay-estuarine system of Goa, central west coast of India"

**Table S1. The COI and Cyt *b* gene sequences**. GenBank accession numbers and sources of the mitochondrial gene sequences (COI and Cyt *b*) of seahorse species used for construction of phylogenetic trees.

| **Species** | **Loci** | **GenBank Accession no.** | **Voucher/field no.** | **Locality** | **Reference** |
| --- | --- | --- | --- | --- | --- |
| *H. mohnikei* | COI | MN595218 | MK1 | Goa, India | Present study |
|  |  | MN595216 | MK2 |  | Present study |
|  |  | MN595217 | MK3 |  | Present study |
|  |  | MK330041.1 | NIO1015/19/  SMkei-05 |  | Present study |
|  | Cyt *b* | MN595213 | MK1 |  | Present study |
|  |  | MN595214 | MK2 |  | Present study |
|  |  | MN595215 | MK3 |  | Present study |
|  |  | MK112274.2 | NIO1015/19/ SMkei-05 |  | Present study |
| *H. mohnikei* | COI | GQ502159.1 | RM 2177i | Japan | Hou et al. (2018) |
|  |  | GQ502158.1 | RM 2177c | Japan | Hou et al. (2018) |
|  |  | GQ502157.1 | RM 2179 | Vietnam | Hou et al. (2018) |
|  | Cyt *b* | AF192689.1 | JAP.346.8 | Japan | Casey et al. (2004) |
|  |  | AF192688.1 | JAP.346.7 | Japan | Casey et al. (2004) |
|  |  | KT731898 | FAKU 135643 | Japan | Han et al. (2019) |
|  |  | KT731899 | FAKU 134645 | Japan | Han et al. (2019) |
|  |  | KT731900.1 | FAKU 135791 | Japan | Han et al. (2019) |
|  |  | KX017612.1 | FAKU 136030 | Japan | Han et al. (2019) |
|  |  | KX017613.1 | FAKU 136031 | Japan | Han et al. (2019) |
|  |  | KX017614. | FAKU 136041 | Japan | Han et al. (2019) |
|  |  | KX017615.1 | FAKU 136042 | Japan | Han et al. (2019) |
|  |  | KX017622.1 | PKU 13175 | Japan | Han et al. (2019) |
|  |  | KX017623.1 | PKU 13176 | Japan | Han et al. (2019) |
|  |  | KX017624.1 | PKU 13177 | Japan | Han et al. (2019) |
|  |  | KX017625.1 | PKU 13178 | Japan | Han et al. (2019) |
|  |  | KX017616.1 | FAKU 136043 | Japan | Han et al. (2019 |
|  |  | KX017617.1 | FAKU 136050 | Japan | Han et al. (2019 |
|  |  | KX017618.1 | FAKU 138439 | Japan | Han et al. (2019 |
|  |  | KX017619.1 | FAKU 139779-1 | Japan | Han et al. (2019 |
|  |  | KX017620.1 | FAKU 139779-2 | Japan | Han et al. (2019 |
|  |  | KX017621.1 | FAKU 139779-3 | Japan | Han et al. (2019 |
|  |  | KT731901.1 | FAKU 137341 | Japan | Han et al. (2019) |
|  |  | KT731902.1 | FAKU 137342 | Japan | Han et al. (2019) |
|  |  | KT731903.1 | FAKU 137343 | Japan | Han et al. (2019 |
|  |  | KT731904.1 | FAKU 137344 | Japan | Han et al. (2019 |
|  |  | KT731905.1 | FAKU 137345 | Japan | Han et al. (2019 |
|  |  | KT731906.1 | FAKU 137346 | Japan | Han et al. (2019 |
|  |  | KT731907.1 | FAKU 137347 | Japan | Han et al. (2019 |
|  |  | KT731908.1 | FAKU 137352, | Japan | Han et al. (2019 |
|  |  | KT731909.1 | FAKU 137353 | Japan | Han et al. (2019 |
|  |  | KX017623.1 | PKU 13176 | Japan | Han et al. (2019) |
|  |  | KT731897.1 | ESFRI 899 | Korea | Han et al. (2019) |
|  |  | KT731937.1 | PKU 11651 | Korea | Han et al. (2019 |
|  |  | KT731910.1 | PKU 6096 | Korea | Han et al. (2019 |
|  |  | KT731911.1 | PKU 6460 | Korea | Han et al. (2019) |
|  |  | KT731935.1 | PKU 11471 | Korea | Han et al. (2019) |
|  |  | KT731919.1 | PKU 11249 | Korea | Han et al. (2019 |
|  |  | KT731920.1 | PKU 11250 | Korea | Han et al. (2019 |
|  |  | KT731921.1 | PKU 11251 | Korea | Han et al. (2019 |
|  |  | KT731922.1 | PKU 11252 | Korea | Han et al. (2019 |
|  |  | KT731931.1 | PKU 11446 | Korea | Han et al. (2019 |
|  |  | KT731936.1 | PKU 11633 | Korea | Han et al. (2019 |
|  |  | KT731930.1 | PKU 11445 | Korea | Han et al. (2019 |
|  |  | KT731923.1 | PKU 11253 | Korea | Han et al. (2019) |
|  |  | KT731924.1 | PKU 11254 | Korea | Han et al. (2019) |
|  |  | KT731925.1 | PKU 11255 | Korea | Han et al. (2019) |
|  |  | KT731926.1 | PKU 11256 | Korea | Han et al. (2019) |
|  |  | KT731928.1 | PKU 11406 | Korea | Han et al. (2019) |
|  |  | KT731913.1 | PKU 11160 | Korea | Han et al. (2019) |
|  |  | KT731914.1 | PKU 11161 | Korea | Han et al (2019) |
|  |  | KT731955.1 | PKU 12411 | Korea | Han et al. (2019) |
|  |  | KT731939.1 | PKU 52866 | Korea | Han et al. (2019) |
|  |  | KT731941.1 | PKU 52870 | Korea | Han et al. (2019) |
|  |  | KT731946.1 | PKU 12007 | Korea | Han et al. (2019) |
|  |  | KT731949.1 | PKU 12412 | Korea | Han et al (2019) |
|  |  | KT731952.1 | PKU 12415 | Korea | Han et al. (2019) |
|  |  | KT731954.1 | PKU 12295 | Korea | Han et al. (2019) |
|  |  | KT731950.1 | PKU 12413 | Korea | Han et al. (2019) |
|  |  | KT731951.1 | PKU 12414 | Korea | Han et al. (2019) |
|  |  | KT731948.1 | PKU 12294 | Korea | Han et al (2019) |
|  |  | KT731945.1 | PKU 54107 | Korea | Han et al (2019) |
|  |  | KT731918.1 | PKU 11248 | Korea | Han et al. (2019) |
|  |  | KT731932.1 | PKU 11447 | Korea | Han et al (2019) |
|  |  | KT731944.1 | PKU 52876 | Korea | Han et al. (2019) |
|  |  | KT731953.1 | PKU 12416 | Korea | Han et al. (2019) |
|  |  | KT731950.1 | PKU 12413 | Korea | Han et al (2017) |
|  |  | KT731947.1 | PKU 12279 | Korea | Han et al. (2019) |
|  |  | KT731933.1 | PKU 11451 | Korea | Han et al. (2019) |
|  |  | KT731915.1 | PKU 11165 | Korea | Han et al (2019) |
|  |  | KT731916.1 | PKU 11166 | Korea | Han et al (2019) |
|  |  | KT731917.1 | PKU 11167 | Korea | Han et al. (2019) |
|  |  | KT731929.1 | PKU 11443 | Korea | Han et al (2019) |
|  |  | KT731934.1 | PKU 11452 | Korea | Han et al (2019) |
|  |  | KT731927.1 | PKU 11288 | Korea | Han et al (2019 |
|  |  | KT731943.1 | PKU 52874 | Korea | Han et al. (2019) |
|  |  | KT731940.1 | PKU 52868 | Korea | Han et al. (2019) |
|  |  | KT731938.1 | PKU 52864 | Korea | Han et al. (2019) |
|  |  | KT731940.1 | PKU 52872 | Korea | Han et al. (2019) |
|  |  | KT731938.1 | PKU 10982 | Korea | Han et al. (2019) |
| *H. mohnikei* | COI | KP140180.1 | SCSMBC001900 | China | Zhang and Lin (2014)* |
|  |  | KP140167.1 | SCSMBC001887 | China | Zhang and Lin (2014)* |
|  |  | KP140168.1 | SCSMBC001888 | China | Zhang and Lin (2014)* |
|  |  | KP140166.1 | SCSMBC001886 | China | Zhang and Lin (2014)* |
|  |  | KP140158.1 | SCSMBC001878 | China | Zhang and Lin (2014)* |
|  |  | KP140157.1 | SCSMBC001877 | China | Zhang and Lin (2014)* |
|  |  | KP140156.1 | SCSMBC001876 | China | Zhang and Lin (2014)* |
|  |  | KP140141.1 | SCSMBC001861 | China | Zhang and Lin (2014)* |
|  |  | KP140153.1 | SCSMBC001873 | China | Zhang and Lin (2014)* |
|  |  | KP140155.1 | SCSMBC001875 | China | Zhang and Lin (2014)* |
|  |  | KP140159.1 | SCSMBC001879 | China | Zhang and Lin (2014)* |
|  |  | KP140160.1 | SCSMBC001880 | China | Zhang and Lin (2014)* |
|  |  | KP140171.1 | SCSMBC001891 | China | Zhang and Lin (2014)* |
|  |  | KP140148.1 | SCSMBC001868 | China | Zhang and Lin (2014)* |
|  |  | KP140179.1 | SCSMBC001899 | China | Zhang and Lin (2014)* |
|  |  | KP140145.1 | SCSMBC001865 | China | Zhang and Lin (2014)* |
|  |  | KP140177.1 | SCSMBC001897 | China | Zhang and Lin (2014)* |
|  |  | KP140164.1 | SCSMBC001884 | China | Zhang and Lin (2014)* |
|  |  | KP140178.1 | SCSMBC001898 | China | Zhang and Lin (2014)* |
|  |  | KP140144.1 | SCSMBC001864 | China | Zhang and Lin (2014)* |
|  |  | KP140170.1 | SCSMBC001890 | China | Zhang and Lin (2014)* |
|  |  | KP140146.1 | SCSMBC001866 | China | Zhang and Lin (2014)* |
|  |  | KP140181.1 | SCSMBC001901 | China | Zhang and Lin (2014)* |
|  |  | KP140147.1 | SCSMBC001867 | China | Zhang and Lin (2014)* |
|  |  | KP140169.1 | SCSMBC001889 | China | Zhang and Lin (2014)* |
|  |  | KP140149.1 | SCSMBC001869 | China | Zhang and Lin (2014)* |
|  |  | KP140137.1 | SCSMBC001857 | China | Zhang and Lin (2014)* |
|  |  | KP140162.1 | SCSMBC001882 | China | Zhang and Lin (2014)* |
|  |  | KP140174.1 | SCSMBC001894 | China | Zhang and Lin (2014)* |
|  |  | KP140173.1 | SCSMBC001893 | China | Zhang and Lin (2014)* |
|  |  | KP140163.1 | SCSMBC001883 | China | Zhang and Lin (2014)* |
|  |  | KP140152.1 | SCSMBC001872 | China | Zhang and Lin (2014)* |
|  |  | KP140140.1 | SCSMBC001860 | China | Zhang and Lin (2014)* |
|  |  | KP140143.1 | SCSMBC001863 | China | Zhang and Lin (2014)* |
|  |  | KP140139.1 | SCSMBC001859 | China | Zhang and Lin (2014)* |
|  |  | KP140150.1 | SCSMBC001870 | China | Zhang and Lin (2014)* |
|  |  | KP140161.1 | SCSMBC001881 | China | Zhang and Lin (2014)* |
|  |  | KP140172.1 | SCSMBC001892 | China | Zhang and Lin (2014)* |
|  |  | KP140154.1 | SCSMBC001874 | China | Zhang and Lin (2014)* |
|  |  | KP140151.1 | SCSMBC001871 | China | Zhang and Lin (2014)* |
|  |  | KP140138.1 | SCSMBC001858 | China | Zhang and Lin (2014)* |
|  |  | KP140165.1 | SCSMBC001885 | China | Zhang and Lin (2014)* |
|  |  | KP140142.1 | SCSMBC001862 | China | Zhang and Lin (2014)* |
|  |  | KP140175.1 | SCSMBC001895 | China | Zhang and Lin (2014)* |
|  | Cyt *b* | KC527556 | HJ1 | China | Zhang et al. (2014) |
|  |  | KC527584.1 | HJ29 | China | Zhang et al. (2014) |
|  |  | KT946937.1 | Haplotype M3 | China | Zhang et al. (2016) |
|  |  | KT946935.1 | Haplotype M2 | China | Zhang et al. (2016) |
|  |  | KT946929.1 | Haplotype M1 | China | Zhang et al. (2016) |
|  |  | KT946932.1 | Haplotype M4 | China | Zhang et al. (2016) |
|  |  | EU179924.1 | moh_TH1 | Thailand | Panithanark (2015)* |
|  |  | EU179923.1 | moh_TH2 | Thailand | Panithanark (2015)* |

*References not cited in the article

Panitharanak T. Phylogeny of Thai seahorses inferred from mitochondrial DNA cytochrome *b* gene. In Proceedings of the Burapha University International Conference 2015 (BUU2015). 2015; pp. 1010–1023). Chonburi: Burapha University.

Zhang Y. Lin Q. DNA barcoding Chinese medicinal fish species in the family, Syngnathidae 2014. Unpublished.
