## Supplementary table 2 for "Morphological and molecular evidence for range extension and first occurrence of the Japanese seahorse, *Hippocampus mohnikei* (Bleeker 1853) in a bay-estuarine system of Goa, central west coast of India"

**Table S2.** **PCA analysis of seahorse species.** Loadings of components matrix and contributions of each axis to total variance ranked as per their coefficients (eigen values > ± 0.30 are included; values in parentheses indicate variance explained by each principal component).

| **Morphological characters** | **Coefficients** |
| --- | --- |
| **Principal component 1 (81%)** |  |
| SL | 0.334 |
| TaL | 0.333 |
| TD4 | 0.329 |
| TD9 | 0.327 |
| HL | 0.327 |
| TrL | 0.325 |
| HD | 0.324 |
| SnL | 0.318 |
| SnD | 0.314 |
| **Principal component 2 (16%)** |  |
| TaR | 0.663 |
| DF | 0.655 |
